# Mixed signatures for subcritical dynamics in rodent hippocampus during sleep and awake epochs

**DOI:** 10.1101/2023.10.30.564597

**Authors:** Pranjal Garg

## Abstract

Neuronal dynamics such as brain criticality have recently been attributed to optimal information processing. Brain criticality attempts to elucidate the collective dynamics of a large number of neurons. It posits that the brain operates near critical to the critical point, although the field is rife with controversies and contrasting evidence. Similar computational capacities are observed during sharp wave ripples in the hippocampus prompting the need to correlate their dynamics. In the current study, the measures of avalanche criticality including neuronal avalanches, branching process, crackling noise relation, and deviation from criticality coefficient and Hurst exponents for long-range temporal correlations in rodent hippocampus during sharp wave ripples are reported. The evidence for mixed subcritical to critical dynamics in the hippocampus and minimal difference between ripple and no ripple times across measured metrics was found. The evidence demonstrates heterogeneity in signatures of criticality among animals and brain areas, indicating the presence of broad-range neuronal dynamics.

---

Criticality, borrowed from statistical physics, is used to study complex brain dynamics, transitioning between disorder (subcritical) and ordered states (supercritical) (1–4). It enhances the brain’s responsiveness to stimuli, information transmission, memory processing, (4–8) and has been observed in various brain regions like the hippocampus during both awake and sleep states (9–11). In the hippocampus, sharp wave ripples (SWRs) have also been observed during both sleep and awake periods, believed to play a role in memory retrieval, consolidation, and planning (12–14). The bulk of SWRs are marked by memory replay and are also a candidate for bidirectional information flow between hippocampus and cortex (15, 16). The intersection of these functions motivated us to explore the criticality in the hippocampus during ripples.

Determination of avalanche criticality requires multiple estimation strategies. To do this, hippocampal local field potential (LFP) data during ripple and no ripple times was analyzed. I primarily hypothesize that criticality and sharp wave ripples are possibly distinct dynamical properties, criticality being more fundamental in nature. It was also found that the hippocampus has distinguishing criticality-related characteristics.

Traditionally, criticality has been determined through the observation of neuronal avalanches, the spontaneous cascades of neural activity characterized by power-law distributions in terms of their size and duration (3).

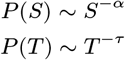

I identified neuronal avalanches during sleep and awake epochs and computed the power law exponents *α* and *τ* (Fig 1. (a)). To discretize and identify the suprathreshold events in LFP variable standard deviations (SD) and bin sizes were applied.

**Figure 1.**
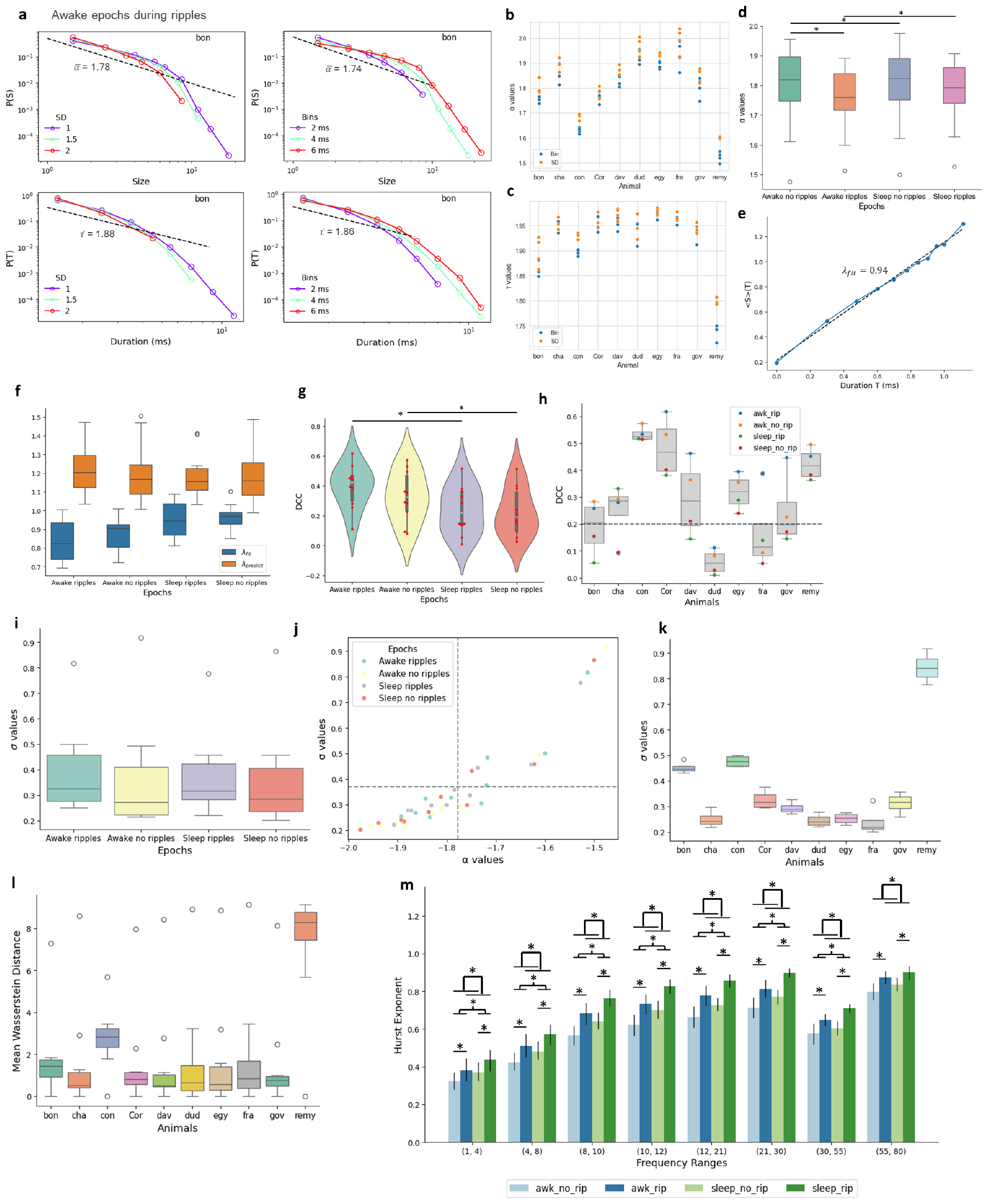
Avalanche, branching, and time metrics for avalanche criticality. **(a)** Power law distribution of avalanches for various thresholds (left panel) and bin sizes (right panels), truncated after a few orders of magnitude (size) and a single order of magnitude (duration). Upper panels represent avalanche size and lower panels represent avalanche durations. **(b)** Avalanche size exponents (*α*) and **(c)** duration exponents (*τ*) for variable threshold and bin sizes. **(d)** Boxplot of *α* values plotted against epochs. (paired t-test, p*<*0.05). **(e)** Line fit (dashed line) with exponent *λ*_*fit*_ to the plot of mean avalanche size *< S >* conditioned on a specific duration T and duration T. **(f)** Boxplot of mean *λ*_*predict*_ and *λ*_*fit*_. **(g)** Violin plots of distance to criticality coefficient (DCC). **(h)** Mean boxplots of DCC for all epochs. **(i)** Mean branching parameter (*σ*) for all animals. **(j)** Mean *α* values plotted against mean *σ* values for all epochs with grey dashed lines indicating the overall mean. **(k)** Mean *σ* values for all epochs. **(l)** Box plot of mean Wasserstein distance between all ten animals. **m** Overall mean Hurst exponents to estimate LRTC across frequency bands. (paired t-test, p*<*0.05).

Before the cut-off marked by the size of avalanches obeying power law, nearly constant power law exponents were found. For varying SD in both avalanche size and lifetime, the power law was preserved, with nearly equivalent power law exponents indicating threshold scale invariance and fractal avalanche organization (17). Similarly, it was preserved for different bin sizes and as expected the absolute value of the exponent increased with increasing bin size, an empirical demonstration of temporal scale-invariance. To estimate the fit of the power law I performed the Kolmogorov Smirnov test. The p-values indicated significant power law distribution for the size of avalanches during sleep and awake states, supporting criticality in the brain.

Original studies showed power law exponents for the size of *α ∼* 3*/*2 and for the duration of *τ ∼* 2. In contrast, variable values of power law exponents were found, although *τ* values were generally higher than *α* values (Fig 1. (b) and Fig. 1 (c)). For different epochs, awake during no ripples (*α* = 1.79 ± 0.14), awake during ripples (*α* = 1.75 ± 0.11), sleep during no ripples (*α* = 1.79 ± 0.13) and sleep during ripples (*α* = 1.77±0.11) I found significant differences in the *α* values during awake and sleep epochs and during ripples and no ripples during awake epoch (Fig. 1 (d)). The findings of values of power law however neither confirm nor rule out criticality.

To evaluate criticality rigorously, I assessed crackling noise scaling relations (18). Scaling theory predicts the distribution between mean sizes conditioned on a specific duration.

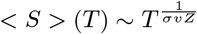

Here, *σ, v*, and *z* are other critical exponents of the system (*σ* is not the branching parameter). In Fig. 1 (e) I have demonstrated the linear fit to determine the exponent 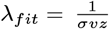. At the critical point, the scaling theory satisfies the relation,

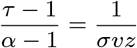

In the current paper, I found that none of the epochs satisfied the scaling relation as shown in Fig. 1 (f), (*λ*_*predict*_ = (*τ −* 1)*/*(*α −* 1)) indicating possible deviation from criticality. To quantify this distance I computed Deviation from Criticality Coefficient defined as (19),

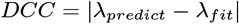

I found a significant difference in DCC (Fig. 1 (g)) between sleep and awake epochs during both ripple and no ripple times, wherein sleep no ripples showed the maximum proximity to criticality. Furthermore, I found variable DCC in different animals. Some animals showed critical dynamics (*DCC <* 0.2) indicating interindividual differences (Fig. 1 (h)). To classify the system further into supercritical, subcritical, and critical states I estimated the branching parameter (*σ*).

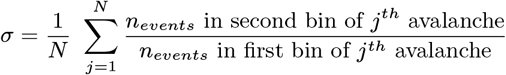

where *N* is the total avalanches in the specific epoch and *n*_*events*_ is the total suprathreshold events in the specific bin. In Fig. 1 (i) I found that across epochs the system was primarily subcritical (*σ <* 0.5). Fig. 1 (j) demonstrates the phase plot of the branching parameter vs avalanche size exponent relationship, with mean values far from *σ* = 1 and *α* = 1.5 suggesting subcriticality in the system. Except for one animal (*remy* in Fig. 1 (b), Fig. 1 (c) and Fig. 1 (k)), all other animals followed this trend. To quantify the differences between individual animals, I calculated the mean Wasserstein distance for the distributions of avalanche size and duration. In Fig. 1 (l), *remy* showed the maximum distance from the rest of the animals (*W* = 7.3 ± 2.68). Although the animals *bon, fra*, and *dud* showed critical dynamics in Fig. 1 (h), they did not have avalanche profiles different from other animals and could indicate limitations of metrics used.

Next, long-range temporal correlations (LRTC), one of the cardinal signs of criticality was considered to estimate the scaling exponent or Hurst exponent (H) (1). H values in the 0.5-1 range indicate non-trivial positive autocorrelation, suggesting the presence of LRTC (20). In Fig. 1 (m) I explored Hurst exponents for sleep and awake epochs for delta band (*<*4 Hz), theta band (4–8 Hz), alpha band (8–12 Hz), beta band (12–30 Hz), and gamma band (30–80 Hz). As expected, exponents were generally larger during sleep than during awake (21). I found the largest exponents for the gamma band across all epochs closely followed by exponents in the beta band. I hypothesize that the observed LRTC might influence the initiation and propagation of SWRs, emphasizing some role of criticality in shaping ripples.

Overall, mixed signatures of subcriticality in the hippocampus in contrast to previous evidence in calcium imaging studies were found (9). Subcriticality is marked by focussed attention, response specificity to stimulus, and stimulus detection among others as compared to supercriticality (5). The dominance of subcriticality over supercriticality is also said to control brain activity from becoming epileptic (22, 23). The mixed signatures and variability in different individuals and previous attempts could possibly be reconciled by the presence of heterogeneous neuronal dynamics across regions (24). While it’s suggested that the hippocampus operates closer to criticality during expensive cognitive tasks, my findings indicate the likelihood of subcritical to near critical dynamics during less demanding tasks (9) in contrast to cortical structures (25). Heterogeneity in results could also be due to experimental and estimation methodology-induced variability. Additionally, non-significant difference between ripples and non-ripple times was observed, suggesting that criticality may not significantly optimize information processing, particularly during ripples in the hippocampus.

## Data Availability

Open access dataset (26) was used to perform the analysis in the current study.

## ACKNOWLEDGMENTS

I am grateful to Benedetta Mariani and Eric Denovellis for helping me access the code for criticality analysis and the data respectively. I would also like to thank Fangchen Zhu for the comments and discussion,

