## Supplementary Information for "Mixed signatures for subcritical dynamics in rodent hippocampus during sleep and awake epochs"

### Supporting Information Text

#### Extended Methods

##### Hippocampus local field potential data and sharp wave ripples

Hippocampus local field potential (LFP) data was obtained from a previously published dataset of recordings from 10 rodents (1). Sharp wave ripples were identified with the accompanied code (2).

##### Neuronal avalanches

The size and duration of the neuronal avalanches were obtained by dividing the LFP into sets of ripple and non-ripple times. In each set the LFP activity was first normalized and then suprathreshold activity was obtained by thresholding them with $-xSD$ (where x = 1, 1.5, 2). These suprathreshold activities were then binned with variable bin sizes (b = 2 ms, 4 ms, 6 ms). An avalanche was identified as a cluster of binned activity bound by empty bins. Before estimating the avalanche metrics, the neuronal avalanches for each animal and each epoch were combined.

##### Power law fit

To estimate the power law exponents for the size and duration of the neuronal avalanches I used the Python package *powerlaw* (3). The goodness of the fit was determined by minimizing the Kolmogorov-Smirnov distance between the synthetic and empirical distribution. The synthetic data (n=1000) was generated using the best-fit power law distribution and the same number of samples as the experimental dataset. The p-values were defined as the deviation from power law by empirical data compared to each synthetic data (4).

##### Wasserstein distance

I computed the Wasserstein distance between the avalanche size and duration distributions of neuronal avalanches using Python's *scipy* (5, 6). Each distribution was weighted by the corresponding counts. To report the final distance I obtained the mean Wasserstein distance for size and duration distributions for each animal.

##### Long range temporal correlations

I estimated long-range temporal correlations with unbiased detrended fluctuation analysis (UDFA) (7) using Python's *fathon* package (8). To apply UDFA, I first filtered LFP with variable frequency bands and then computed the amplitude envelope.
